# Transposable elements acquire time- and sex-specific transcriptional and epigenetic signatures along mouse fetal gonad development

**DOI:** 10.1101/2023.10.23.563600

**Authors:** Isabelle Stévant, Nitzan Gonen, Francis Poulat

## Abstract

Gonadal sex determination in mice is a complex and dynamic process, crucial for the development of functional reproductive organs. The expression of genes involved in this process is regulated by a variety of genetic and epigenetic mechanisms. Recently, there has been increasing evidence that transposable elements (TEs), which are a class of mobile genetic elements, play a significant role in regulating gene expression during embryogenesis and organ development. In this study, we aimed to investigate the involvement of TEs in the regulation of gene expression during mouse embryonic gonadal development. Through bioinformatic analysis, we aimed to identify and characterize specific TEs acting as regulatory elements for sex-specific genes, as well as their potential mechanisms of regulation. We identified TE loci expressed in a time- and sex-specific manner along fetal gonad development that correlate positively and negatively with nearby gene expression, suggesting that their expression is integrated to the gonadal regulatory network. Moreover, chromatin accessibility and histone post-transcriptional modification analyses in the differentiating supporting cells revealed that TEs are acquiring sex-specific signature for promoter-, enhancer-, and silencer-like elements with some of them being proximal to critical sex determining genes. Altogether, our study introduces TEs as new potential players of the gene regulatory network controlling gonadal development in mammals.

## Introduction

Sex determination is the developmental process by which individuals acquire the necessary organs for sexual reproduction. In vertebrates, it starts with the differentiation of the bipotential gonadal primordium into ovaries or testes. In mice, the bipotential gonadal forms around embryonic day (E)10.0 at the ventral surface of the intermediate mesoderm and is composed of multipotent somatic cells and primordial germ cells (Mayere et al. 2022, Neirijnck et al. 2023). At this stage, the XX and XY somatic cells display no autosomal sexual dimorphism at the transcriptional (Stevant et al. 2019) and chromatin levels (Garcia-Moreno, Futtner, et al. 2019). The supporting cell lineage is the first to operate sex fate decision at E11.5 by differentiating as either Sertoli cells or pre-granulosa cells depending on the presence or absence of the Y chromosome (Albrecht and Eicher 2001, Chassot et al. 2008). Once specified, Sertoli and pre-granulosa cells instruct other somatic progenitors and the primordial germ cell to commit and differentiate towards the testicular or ovarian cell fates. In XX gonads, the canonical WNT pathway (WNT4/RSPO1/β-catenin) and the transcription factor (TF) FOXL2 induce the pre-granulosa cells differentiation (review:(Greenfield 2021)). In XY gonads, sex determination is governed by the *Sry* gene located on the Y chromosome (Koopman et al. 1991, Sinclair et al. 1990) that activates at E11.5 the expression of the transcription factor SOX9, (Gonen et al. 2018) which in turn controls Sertoli cells differentiation (review: (Vining et al. 2021)). In both sexes, the action of master transcription factors induces the activity of sex-specific regulatory networks, controlling the expression and repression of downstream pro-testicular or pro-ovarian genes. Many levels of regulation are expected to play a role in controlling this delicate gene regulatory network. Indeed, it has been shown that prior to sex determination, sex-specific genes carry a bivalent histone mark signature with both active (H3K4me3) and repressive (H3K27me3) marks. As sex differentiation progresses, these genes lose one of these marks and acquire sex-specific expression (Garcia-Moreno, Lin, et al. 2019). Furthermore, many sex specific intergenic loci acquire deposition of H3K27Ac, which characterizes active enhancers (Garcia-Moreno, Futtner, et al. 2019). With two alternative outcomes, the study of sex determination offers an attractive model for studying the epigenetic events involved in cell fate decisions.

We have previously shown that the versatile nuclear-scaffold protein TRIM28 directly interacts and co-localizes with SOX9 on the chromatin of fetal Sertoli cells (Rahmoun et al. 2017). We also showed that TRIM28 protects adult ovaries against granulosa-to-Sertoli cell reprogramming through it SUMO-E3 ligase activity (Rossitto et al. 2022). Interestingly, in addition to its role on gene regulation as transcriptional activator or repressor, TRIM28 is a master regulator of transposable element (TE) silencing in somatic cells (Rowe et al. 2010). In mammalian genomes, nearly half of the DNA consists of TEs. TEs are divided into two classes depending on the mechanism by which they transpose. Class I TEs are retrotransposons that propagate via a “copy-paste” mechanism by using an intermediate RNA molecule that is reversed transcribed as DNA before re-insertion in the host genome. Class II TEs are DNA transposons propagating via a “cut-and-paste” mechanism. They encode a transposase that excises their flanking inverted terminal repeats and inserts them somewhere else in the host genome. Repression of somatic or germinal TE expression is crucial to block the genetic instability that their uncontrolled expression would induce (Hedges and Deininger 2007). Therefore, most of the TE loci are silenced by H3K9me3 or CpG-methylation in somatic cells and by the piRNAs machinery in male germinal lineage (Zhou et al. 2022). In somatic cells, TRIM28 negatively regulates retrotransposons upon its interaction with KRAB-containing zinc-finger proteins. Consequently, TRIM28 recruits the histone methylase SETDB1 that deposits the heterochromatin mark H3K9me3 contributing to epigenetic retrotransposon silencing (review see: (Randolph et al. 2022)).

Apart from the potential genomic stability threat they represent, some TEs have become intrinsic part of the genome along vertebrate evolution and are drivers of genetic innovations, notably in sex determination and reproduction (review: (Dechaud et al. 2019)). For instance, TE insertion in the sablefish and medaka generated allelic diversification leading to the creation of a new master sex-determining gene (Herpin et al. 2021, Schartl et al. 2018). More recently, a second exon of the *Sry* gene was identified in mice. This cryptic exon originates from the insertion of four retrotransposons. While the short SRY isoform carries a C-terminal degron which renders the protein unstable, the long isoform containing the cryptic exon is degron-free and thus more stable (Miyawaki et al. 2020). Furthermore, co-option and domestication have repurposed TEs for the beneficial of their host and contributed to novel regulatory mechanisms. We distinguish three different mechanisms by which TEs influence gene expression. First, TE are involved in chromatin organization. TEs containing CTCF (CCCT-C binding factor) motifs were found to be directly involved in the formation of topologically associated domains (TADs) and long-range enhancer-promoter interactions (see review (Lawson et al. 2023)). Second, TE sequences are rich in motifs for lineage-specific transcription factors and can be co-opted as *cis*-regulatory elements. As an example, endogenous retroviruses (ERVs) were shown to be highly enriched in specie-specific placenta development enhancers (Chuong et al. 2013). Finally, TE-derived transcripts were shown to influence gene expression by mechanisms that are still poorly understood. In mice, the transition from the 2-cell stage and development progression to the blastocyst stage appear to depend on LINE-1 expression (Jachowicz et al. 2017, Percharde et al. 2018). TE expression was also shown to be involved in adult neurogenesis, neuronal pathologies, and cancer (for review see respectively (Jansz and Faulkner 2021, Jonsson et al. 2020, Richardson et al. 2014)).

In this work, we aim to characterize TE expression and chromatin landscape as fetal gonads specify as testis or ovary in order to establish the groundwork for further functional studies of TEs in mouse gonadal development. We first identified that common TEs were dysregulated in adult ovaries from *Trim28* KO mice (Rossitto, et al. 2022) and ovarian *Dmrt1* overexpression (Lindeman et al. 2015), two models presenting granulosa-to-Sertoli transdifferentiation, suggesting a sex-specific signature of TE expression in mouse. Hence, we reanalyzed transcriptomic and epigenomic data from embryonic gonads and found that indeed major expression of TE loci is detectable in both sexes with temporal- and sex-dependent variation. We observed that a significant proportion of open chromatin regions contain TE sequences associated to active or repressive histone marks and having enrichment for DNA motifs recognized by transcription factors involved in sex determination. Therefore, our findings suggest that TEs may play a role at several levels during mammalian sex determination.

## Results

### Quantification of TE expression in mouse developing gonads

We first reanalyzed RNA-seq data produced from *Trim28* knockout in the fetal pre-granulosa cells that induce a granulosa-to-Sertoli cell reprograming in adult ovaries (Rossitto, et al. 2022), and measured the change in TE expression. Due to their highly repetitive nature and low RNA abundance, evaluation of TE expression requires the use of dedicated tools. TE-specific mapping software re-attributes the sequencing reads, habitually discarded from regular RNA-seq, to TE family or TE loci, allowing the quantification of their expression levels (review: (Lanciano and Cristofari 2020)). For that aim, we used the SQuIRE suite of tools which is based on an expectation-maximization (EM) algorithm to map RNA-seq data and quantify individual TE copy expression (Yang et al. 2019). We found that 8,110 TEs were up-regulated upon cell reprograming, while 2,102 were down regulated (**Sup. Figure 1A**). We predict that the TE dysregulation is mainly driven by the absence of the TE master regulator TRIM28, but we hypothesized that TEs could also be dysregulated by the change of cell identity. To verify this hypothesis, we also reanalyzed RNA-seq data stemming from the reprograming of adult granulosa to Sertoli cells upon *Dmrt1* forced expression (Lindeman, et al. 2015). While DMRT1 has no identified role in TE expression regulation, we found that 4,099 TEs were differentially expressed upon cell reprogramming (**Sup. Figure 1B**). We compared the differentially expressed TEs in the *Trim28* knockout and the *Dmrt1*-induced cell reprograming and found 935 TEs that are commonly dysregulated (**Sup. Figure 1C**), supporting the idea of a sex-specific signature of TE expression in the mouse gonads.

We next sought to evaluate the involvement of TE-derived RNAs in the dynamics of gonadal transcriptomes along sex determination. For that aim, we measured TE RNAs at a locus-specific level in mice using bulk RNA-seq data set of whole mouse gonads from E10.5, E11.5, E12.5 and E13.5 of both sexes (Zhao et al. 2018). We were able to detect between 61,000 and 101,000 expressed TE loci across all the embryonic stages and sexes (**Sup. Figure 2A**). TEs are interspersed throughout the mouse genome and can be located with genes and as such incorporated within gene transcripts. To discriminate TE transcripts induced by their own promoters from passive TE co-transcription with genes, we classified TEs with respect to their environment as done by others (Chang et al. 2022). We considered expressed TEs found in intergenic regions or inside genes that are not transcribed as “self-expressed” since their expression is likely to be driven by their own promoters, while TEs located within expressed genes are qualified as “gene-dependent” since their transcripts are likely to be part of their host gene RNAs (**Figure 1A**). Although TEs are relatively rare in gene bodies (Kapusta et al. 2013), we found that up to 83% of the detected TEs were located with expressed genes. On the other hand, self-expressed TEs constitute around 17% of the detected TEs, which represents between 12,672 and 17,517 TE loci transcribed independently from genes across stages and sexes (**Figure 1B**). In the present study, we focused our interest on the self-expressed TEs as part of autonomously transcribed RNAs susceptible to take part in the gonadal sex differentiation genetic program. As TEs expression has been observed in male germ cells (Liu et al. 2014), we verified if the self-expressed TEs we detected can be expressed by gonadal somatic cells. To this aim, we reanalyzed the only available bulk RNA-seq data from purified somatic cells that was performed at E11.5 in XX and XY gonads (Miyawaki, et al. 2020). We detected 8,696 and 7,479 common self-expressed TEs between whole gonad and somatic cell in XX and XY respectively, which represent almost half of the self-expressed TEs we detected in whole gonads at E11.5 for both sexes (**Sup. Figure 2C**). These results show that the self-expressed TEs we detected in the whole gonads are transcribed by both/either the germ cells and the somatic cells of the gonads.

**Figure 1.**
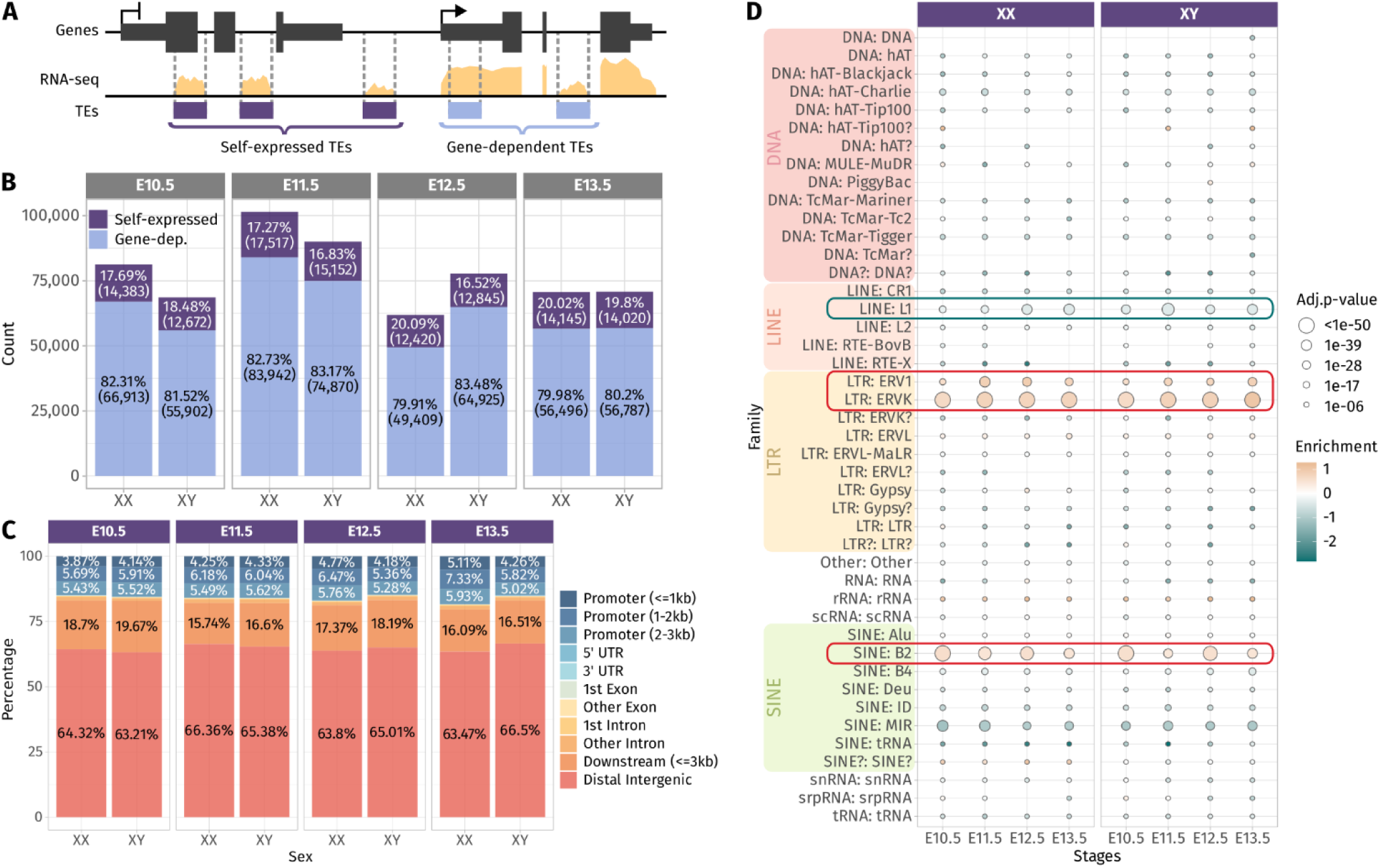
Expressed TEs in the developing gonads. **A)** Classification of the TEs as “self-expressed” when they are found in intergenic regions or within a gene that is not expressed; or as “gene-dependent” when they are found within a transcribed gene. **B)** Number of Self-expressed and Gene-dependent TEs per sex and stage. **C)** Repartition of the self-expressed TEs in the genome for each sex and stage. **D)** Enrichment test of the self-expressed TE families compared to their distribution in the mouse genome. Positively enriched TE families are colored in orange, while the under-represented families are colored in green. Enrichment score is represented as the log2(odds ratio) from the Fisher-exact test. The size of the dots reflects the statistical power of their enrichment. Only enrichment with adj.p-value<0.05 are shown.

We asked whether the self-expressed TEs were located in close proximity of genes, and in particular to promoter regions and 3’ end regions, as TEs have been described to produce alternative promoters, TSSs (transcription start sites) and TTS (transcription termination sites) of genes (Conley and Jordan 2012, Miao et al. 2020). We found that around 15% of the self-expressed TEs were located in gene promoter regions up to 3 kb from the gene TSS, and 20% of them were found in the 3 kb downstream of the 3’ end of the genes. The vast majority of the self-expressed TEs (65%) were distal intergenic located more than 3 kb away from genes (**Figure 1C**). We then performed an enrichment analysis to detect over- or under-represented family of self-expressed TEs compared to the distribution of TE families along the mouse genome (**Figure 1D**). The enrichment test revealed that the DNA families were under-represented among the self-expressed TEs, which is expected as they are mostly inactive and only exist as relics of anciently active elements (Platt et al. 2018). We also notice that the LINE family, and in particular the LINE1 subfamily, was under-represented. LINE1 represents 24.4% of the mouse TEs and ∼1,000 copies are potentially capable of active retrotransposition (Goodier et al. 2001). While LINE1 are highly expressed in the mouse preimplantation embryos (Jachowicz, et al. 2017), fewer copies than expected (8% less) were found transcriptionally active in the developing gonads (**Figure 1D** and **Sup. Figure 2C**). Conversely, we observed a positive enrichment for the ERV1 and ERVK LTRs as well as the B2 SINE non-autonomous retrotransposons.

Overall, we found that TEs are broadly expressed along gonadal development and a large proportion of the detected TEs are constituent part of genes. The detected self-expressed TE loci from the whole gonads are potentially transcribed in both the somatic and the germ cell compartment of the gonads. Self-expressed TEs are mostly found in distal intergenic regions and are enriched in ERVs and SINE B2 retrotransposon families.

### Mouse gonads exhibit gradual sex-specific expression of TEs

After having depicted the TE expression landscape in the developing gonads, we undertook to identify if the self-expressed TE loci change in expression along gonadal differentiation and if they present sexual dimorphism. We performed differential expression analysis, firstly across embryonic stages in both sexes separately, and secondly between XX and XY gonads for each stage. Regarding TEs expression throughout gonadal development, we identified 542 and 324 TEs presenting a dynamic expression along gonadal specification as ovary or testis respectively (**Figure 2A** and **Sup. Data 1**). Dynamically expressed TEs were classified in seven groups (numbered 1 to 7) according to their expression profile. We observed that TEs are expressed in successive waves along gonadal development, with some TEs overexpressed at only one stage (groups 1, 3 and 7 in XX and XY), while other TEs overexpressed during several stages (groups 2, 4 to 6 in XX and groups 2 and 5 XY gonads). Concerning sexual dimorphism, only 6 and 12 self-expressed TEs were detected as differentially expressed between XX and XY at E10.5 and E11.5 respectively (**Figure 2B** and **Sup. Data 2**). Among them, 11 out of the 15 TEs found overexpressed in XY at E10.5 and E11.5 were located on the Y chromosome. These results show that autosomal TEs are not expressed in a sexually dimorphic way in the gonadal cells prior to sex determination. From E12.5, we observed an increase of sexually dimorphic self-expressed TE loci, with 200 and 421 loci differentially expressed between XX and XY gonads at E12.5 and E13.5, respectively. This progression of TE expression perfectly copy what was previously observed concerning the dynamics of expression of the protein-coding genes (Jameson et al. 2012, Nef et al. 2005, Stevant, et al. 2019, Zhao, et al. 2018), except that we observed more overexpressed TEs in XX than XY from E12.5 onward.

**Figure 2.**
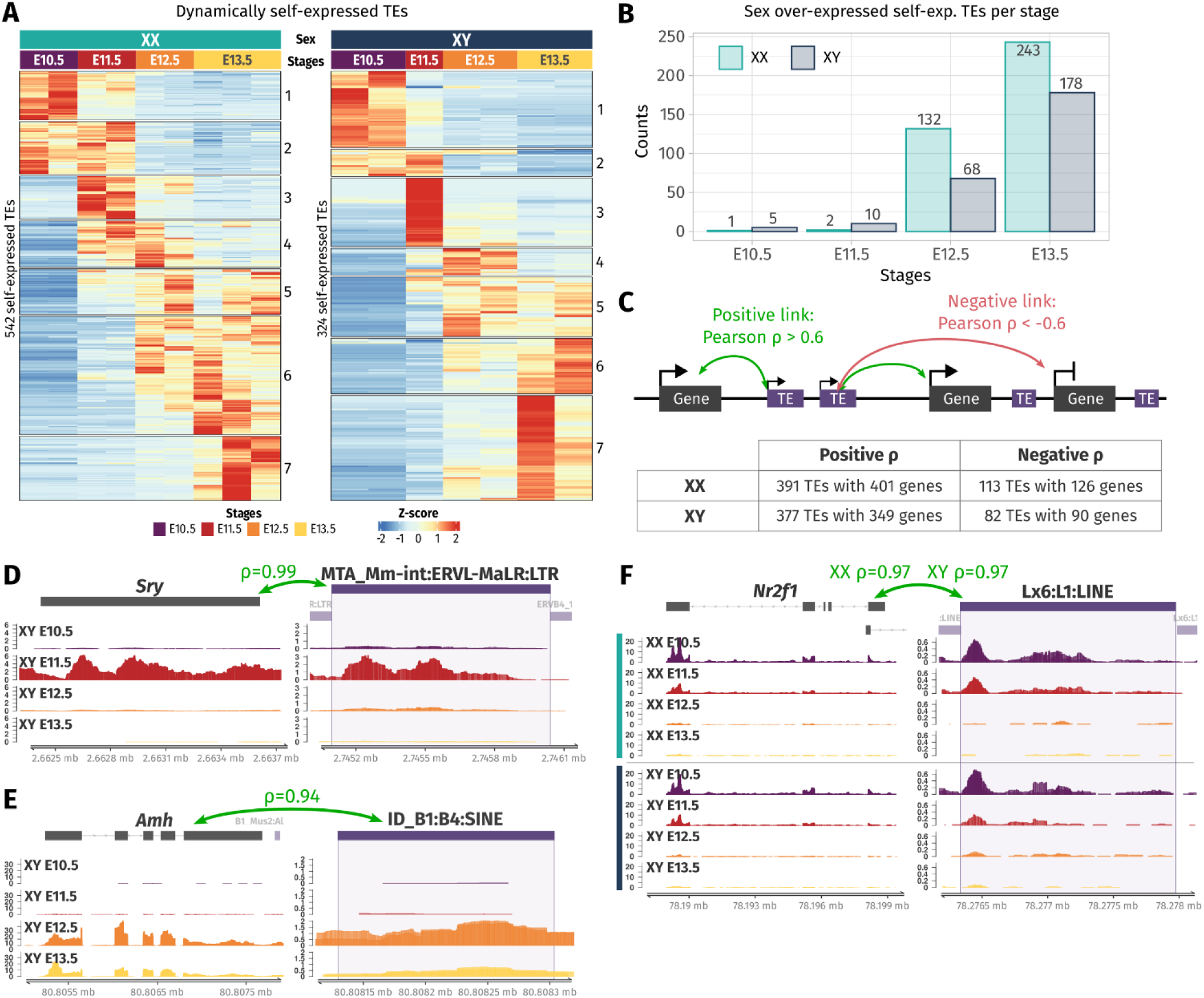
Dynamic and sex-specific TEs along gonadal development. **A)** Heatmap showing the self-expressed TEs that are differentially expressed among the different embryonic stages in XX (left) and XY (right) gonads. The expression is normalized by z-score. **B)** Number of over-expressed self-expressed TEs between sexes for each stage. **C)** Schematic representation describing the strategy to find correlation between TE and their nearby genes. A TE is correlated to a gene in its 100 kb flanking region if the correlation coefficient is above 0.6 (positive correlation) or below −0.6 (negative correlation). The table shows the number of correlating TE and genes found by the analysis. **D-F)** Genomic tracks of the RNA-seq signal of three TE-gene positive correlation examples.

As TE expression can influence the expression level of their nearby genes, we sought to investigate if the expression level of the identified differentially expressed TEs both in time and sex correlates with proximal gene expression. Positive TE-gene correlation can happen for several reasons. The TE transcript can influence the expression of the gene as a non-coding RNA or enhancer RNA (Liang et al. 2023), the TE can act as a promoter or an alternative 3’ UTR for the gene, the gene can be silenced as a collateral effect of the TE silencing, or the TE expression is activated by the same *cis* or *trans* regulatory elements as the gene. On the contrary, negative TE-gene correlation can suggest a negative control of the nearby gene by the TE transcript. As most of the self-expressed TEs were found in a window of 100 kb of a gene TSS (**Sup. Figure 2D**), we looked at the genes present in a 100 kb window around each differentially expressed TE in time and sex and we calculated the Pearson correlation between the gene and the TE expression among embryonic stages and sex (**Figure 2C**). In XX gonads, 391 TE loci were positively correlating with 401 genes, and 113 TEs were negatively correlating with 126 genes. In XY gonads, 377 TEs were positively correlating with 349 genes, while 82 TEs negatively correlated with 90 genes (**Sup. Data 3**). Interestingly, among them we found an LTR ERVL-MaLR retrotransposon located 81 kb upstream of *Sry* expressed with an extremely strong correlation with *Sry* gene expression (ρ=0.99, **Figure 2D**). This ERVL-MaLR is separated from *Sry* by one unexpressed gene (*H2al2b*), demonstrating its expression was produced by a distinct transcript, conversely to the recently identified *Sry* cryptic exon (Miyawaki, et al. 2020). Interestingly, the expression of this TE was also retrieved in RNA-seq data from XY somatic cells purified at E11.5 (Miyawaki, et al. 2020), strongly suggesting that it is expressed in the same cell lineage as *Sry*. We also found a SINE B4 retrotransposon located 2.9 kb downstream of the *Amh* gene with a correlation coefficient of 0.94 (**Figure 2E**). This specific transposon was not expressed in XX gonads where *Amh* is silenced. Its proximity with the *Amh* 3’ UTR and the presence of other expressed TEs in the vicinity can suggest the transcript SINE B4 might be part of an alternative 3’ UTR. Finally, we found five LINE1 TEs positively correlating with the *Nr2f1* gene, also known as *Coup-TFI*, which is expressed in the gonadal somatic cells prior to sex determination in both sexes (Stevant, et al. 2019). *Nr2f1* is a pleiotropic gene able to both activate and repress target gene expression by interacting with chromatin remodelers. It has been described as a driver of cell fate decision in mouse and human brain development (Bertacchi et al. 2019). The LINE1 shown in Figure 2F is part of the second exon of the *lnc-Nr2f1*, (also referred as *A830082K12Rik*) which is located directly upstream of *Nr2f1* on the opposite strand and is conserved between mouse and human (Ang et al. 2019). Down regulation of the *lnc-Nr2f1* in mice causes neural crest cell premature differentiation as glial cells (Bergeron et al. 2016). Both *Nr2f1* and *lnc-Nr2f1* expression prior to gonadal cell differentiation suggest a role in the establishment of the bipotential progenitor cell identity.

To summarize, we found that self-expressed TEs are dynamically expressed and acquire sexual dimorphism along gonadal differentiation. Dynamic and sexually dimorphic TE expression correlates with nearby genes, some of which having a pivotal role in gonadal development. Although we cannot determinate with the present data whether the TE expression precedes or not their nearby gene transcription, this suggests that TEs are a whole part of the genetic program driving gonadal specification.

### TE loci represent one third of open chromatin regions in embryonic supporting cells

Independently to their transcription, TE sequences can influence gene expression by acting as *cis*-regulatory elements including promoters or enhancer by providing a large repository of potential transcription factor binding sites. A comparative study in mouse developing tissues showed that 21% of the open chromatin regions were associated with TEs and half of them were tissue-specific, suggesting an active role in mouse organogenesis (Miao, et al. 2020). We asked whether the chromatin around TE loci is also specifically opening while the bipotential gonads differentiate as ovary or testis. We took advantage of the pioneer epigenetic investigations on mouse embryonic gonads performed by Danielle Maatouk and colleagues (Garcia-Moreno, Futtner, et al. 2019, Garcia-Moreno, Lin, et al. 2019) and re-analyzed the ATAC-seq data of purified gonadal somatic cells at E10.5 and supporting cells at E13.5 in both sexes to identify nucleosome depleted TEs. In total, we identified between 57,014 and 88,312 accessible chromatin regions. Despite inherent mappability issues due to TE repetitive sequences nature (*i.e.* multi-mapped reads are filtered out from the ATAC-seq data), we found that 30% to 39% of the ATAC-seq peaks were overlapping TE loci across sexes and stages (**Figure 3A**). In most cases, more than half of the length of the TE sequences was covered by an ATAC-peak, showing that most of the TE sequence is accessible for potential regulatory factor binding (**Sup. Figure 3**).

**Figure 3.**
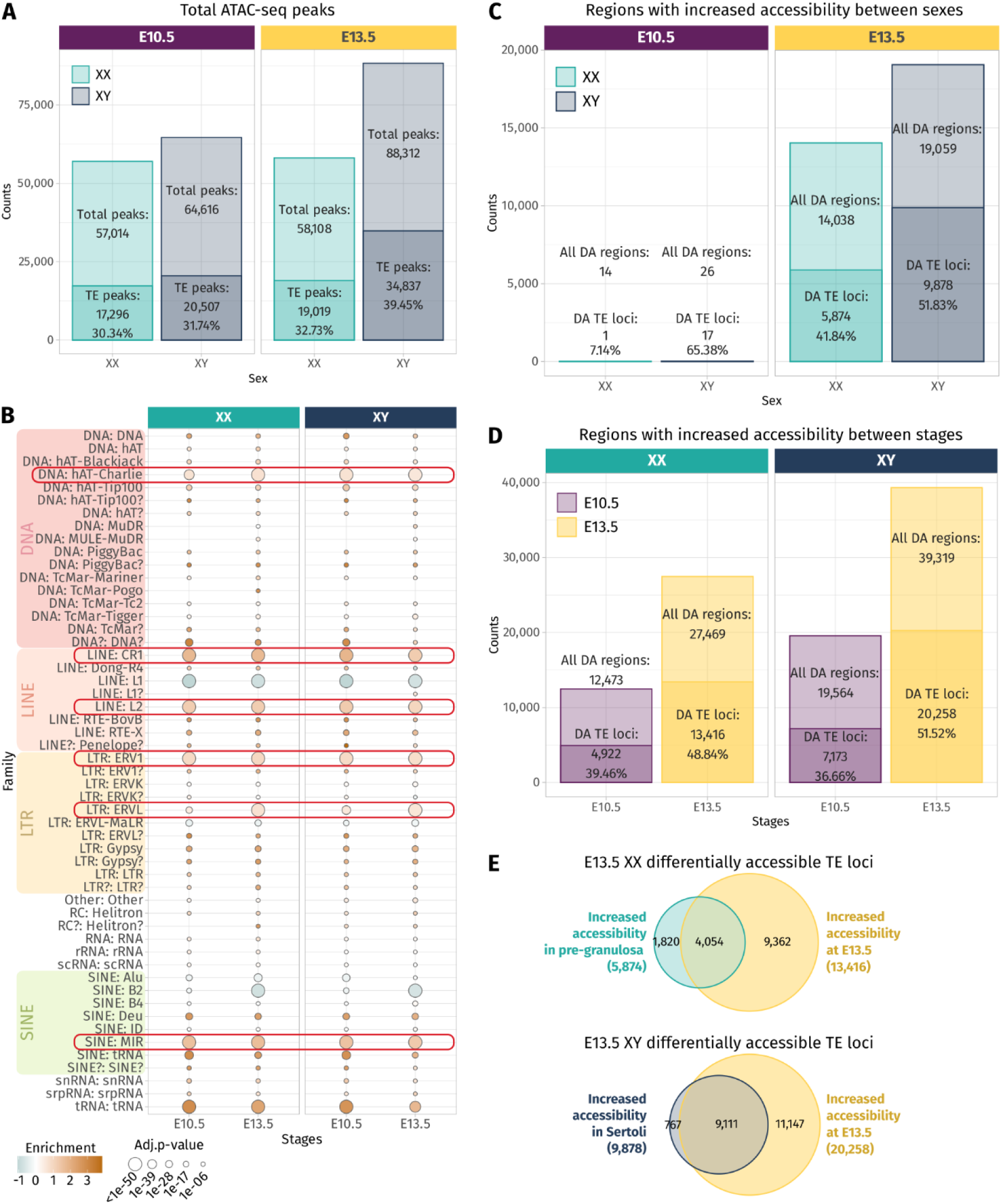
Chromatin accessibility landscape of TE loci during gonadal development. **A)** Number of ATAC-seq peaks detected in total and overlapping TE loci at E10.5 and E13.5 in XX and XY gonadal progenitor and supporting cells. **B)** Family enrichment analysis of the TE overlapping ATAC-seq peaks. Positively enriched TE families are colored in orange, while the under-represented families are colored in green. Enrichment score is represented as the log2(odds ratio) from the Fisher-exact test. The size of the dots reflects the statistical power of their enrichment. Only enrichment with adj.p-value<0.05 are shown. **C-D)** Statistically differentially accessible (DA) regions in sex and time. The number and the percentage of accessible regions and TE containing loci showing an increase of accessibility between sex and time are indicated. **E)** Overlapping of the accessible TE loci that increase in accessibility between sex at E13.5 in either Sertoli and pre-granulosa cells (turquoise and dark blue, respectively) with the accessible TE loci that gain accessibility at E13.5 compared to E10.5 (yellow).

Enrichment test shows that many TE families were overrepresented in the open chromatin regions compared to their global proportion in the genome (**Figure 3B**), in particular the DNA hAT-Charlie, the LINE CR1 and L2, the LTR ERV1, ERVL, and the SINE MIR. These six TE families were also found enriched in TE-derived enhancer-like sequences identified in different human tissue cell lines (Cao et al. 2019), suggesting that they are not necessarily specific to our model and that some TE families are more potent to contribute as *cis*-regulatory elements than others across mammals. We also noticed that the SINE B2 family, which was enriched among the transcribed TE loci in the whole developing gonad (**Figure 1D**), is under-represented among the open chromatin regions in the supporting cells for both sexes. As we expect transcribed TE loci to be accessible, we can speculate that the B2 family is not preferentially transcribed in the somatic cells but the germ cells.

We next performed differential chromatin accessibility analysis between sexes and stages in order to identify the regions that gain in accessibility while gonadal progenitor cells differentiate as either pre-granulosa or Sertoli cells. As previously reported, almost no differences in open chromatin regions between XX and XY were found at E10.5 when the cells are multipotent (Garcia-Moreno, Futtner, et al. 2019). In contrast, at E13.5 we observed 14,038 genomic regions with increased accessibility in pre-granulosa cells compared to Sertoli, and 41.84% of them are overlapping with TE loci. On the other hand, 19,059 genomic regions showed increased accessibility in Sertoli cells compared to pre-granulosa, and 51.83% of them were containing TE loci (**Figure 3C**). We also looked at the genomic regions that increased in accessibility in a time-specific manner in both sexes and found that 27,469 and 39,319 regions were more accessible at E13.5 in pre-granulosa and Sertoli cells, respectively, compared to the E10.5 progenitor cells (**Figure 3D**). Among them, 48.84% and 51.52% were overlapping TE loci in pre-granulosa and Sertoli cells respectively. Finally, we found that 30% and 45% of the TE loci increasing in accessibility at E13.5 in pre-granulosa and Sertoli cells, respectively, were also sexually dimorphic (**Figure 3E**).

Altogether, these results show that TE loci are increasing in accessibility in a time- and sex-specific manner as supporting cells differentiate as pre-granulosa or Sertoli cells suggesting they are participating in the acquisition of the supporting cell type identity as active *cis*-regulatory regions.

### TEs acquire sex-specific enhancer- and promoter-like chromatin landscape as supporting cells differentiate

To reveal the potential roles of TE sequences that become accessible in pre-granulosa and Sertoli cells in a time- and sex-specific manner, we explored the histone post-transcriptional modifications present in their vicinity. We reanalyzed ChIP-seq data for H3K4me3 (marker of promoters), H3K27ac (marker of active promoters and enhancers), and H3K27me3 (marker of silencers and transcriptionally silenced genes by PRC2) performed on purified gonadal somatic cells at E10.5 and supporting cells at E13.5 in both sexes(Garcia-Moreno, Futtner, et al. 2019, Garcia-Moreno, Lin, et al. 2019). We classified the accessible TEs according to the combination of histone mark peaks overlapping with the TE sequences +/-200 bp to have an overview of the chromatin landscape at a one nucleosome resolution (**Sup. Data4**). Finally, we represented the ATAC-seq and the different ChIP-seq signals on the accessible TEs and up to 2 kb around them as juxtaposed heatmaps for the pre-granulosa and the Sertoli cells (**Figure 4A-B**).

**Figure 4.**
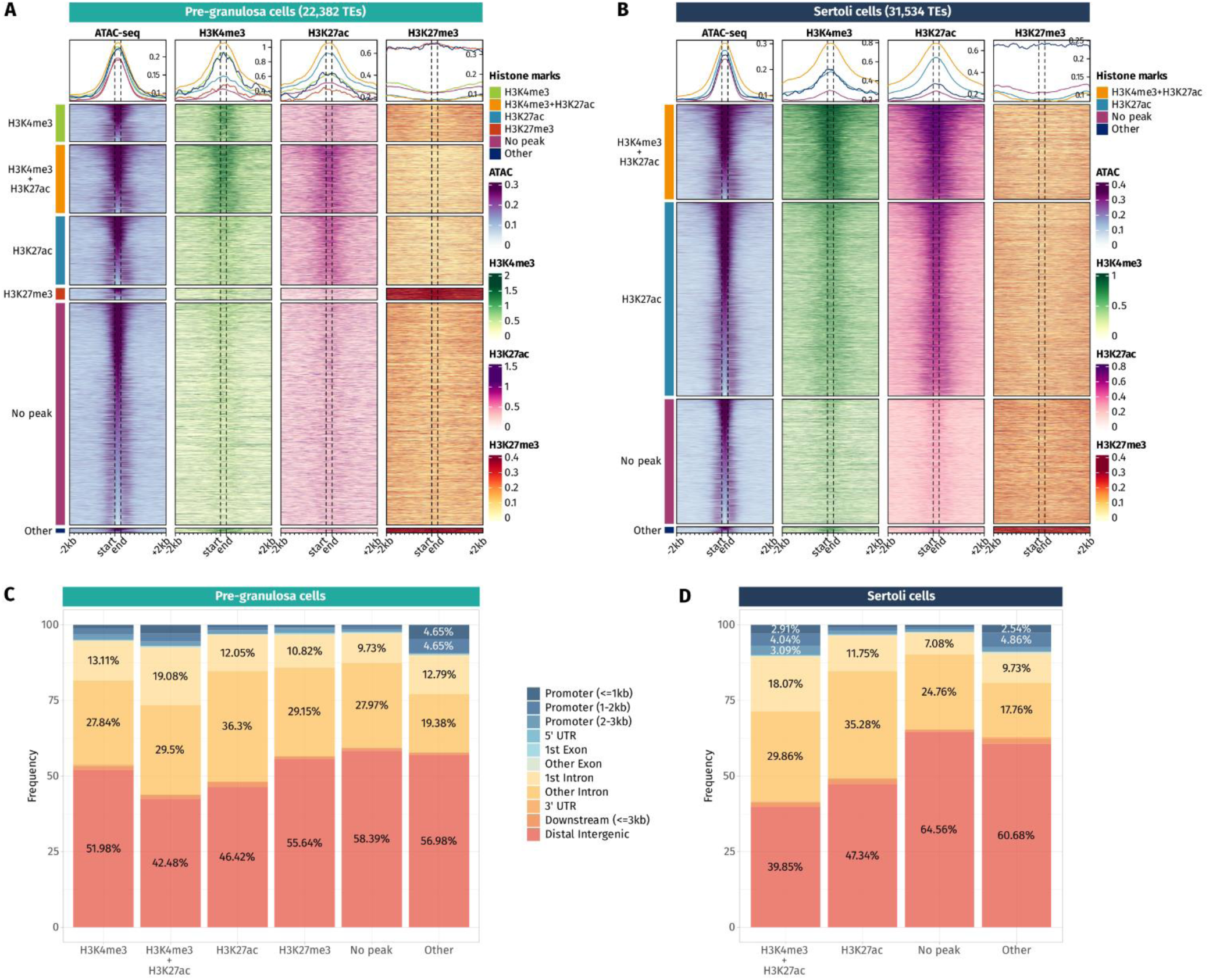
Epigenetic landscape of pre-granulosa and Sertoli cell accessible TE loci. **A-B)** Density plots and heatmaps representing the ATAC-seq and the ChIP-seq signals for histone marks H3K4me3, H3K27ac, and H3K27me3 around the TE loci that gain accessibility in a sex- or time-specific manner in pre-granulosa and Sertoli cells respectively. Accessible TEs were grouped according to the histone marks they display. The groups of histone marks with fewer than 500 TEs were grouped in the “Other” section as the groups were not clearly visible on the final heatmap. **C-D)** Genomic location for each chromatin landscape group of the TE loci that gains accessibility in a sex- or time-specific manner in pre-granulosa and Sertoli cells, respectively.

In pre-granulosa cells, TEs were classified into six histone combination groups (**Figure 4A**). The first two groups contain open TEs displaying histone marks specific for promoters (H3K4me3 and H3K4me3 + H3K27ac). They represent 25.6% of the total of the pre-granulosa enriched accessible TEs (5,727 out of 22,328 TEs). Surprisingly, only 6.3% of them (360 TEs) were found in a gene promoter region (up to 3 kb upstream of a gene TSS) (**Figure 4C**). We looked at the closest genes of the 5,727 H3K4me3 and H3K4me3 + H3K27ac TEs and found critical pre-granulosa cell markers such as *Wnt4*, *Fst*, *Runx1*, *Lef1*, and *Kitl* (**Sup. Data 4**). GO term analysis confirmed statistical enrichments of terms related to reproductive development and Wnt signaling (**Sup. Data 5**). The third group of pre-granulosa enriched accessible TEs displays the H3K27ac histone mark which suggests they are potential enhancers (3,725 out of 22,382). The proximal genes were enriched for the biological process GO term of sex differentiation and included *Gata4*, *Cited2*, *Kitl*, and *Lhcgr* (**Sup. Data 5**). Next, the fourth group contains accessible TEs with the H3K27me3 histone mark that labels silenced genes as well as silencer elements (638 out of 22,328). Most of them are found in distal intergenic regions (**Figure 4C**). Interestingly, we found two H3K27me3 TEs (a SINE B4 and a DNA TcMar-Tigger) in close proximity with *Sox9,* which are located within TESCO (testis specific enhancer of *Sox9* core) (Sekido and Lovell-Badge 2008). We also found a negative regulator of the Wnt pathway, *Sfrp1*, that is expressed in the gonadal progenitor cells but silenced upon the differentiation of the supporting cell lineage of both sexes (Stevant, et al. 2019 17133, Warr et al. 2009). Finally, 53.8% of the pre-granulosa accessible TEs were not overlapping with any of the investigated histone marks. The proximal genes of these TEs were enriched for female sex differentiation and regulation of canonical Wnt signaling pathway GO terms (**Sup. Data 5**).

In Sertoli cells, TEs were classified into four histone combination groups (**Figure 4B**). The first group is composed of TEs displaying promoter-like histone marks H3K4me3 + H3K27ac (7,124 out of 31,534 TEs). As observed in pre-granulosa, only 10% (715 TEs) of them are found in the promoter region of genes (**Figure 4D** and **Sup. Data 4**). The second group shows TEs with enhancer histone mark H3K27ac (17,570 out of 31,534). Like for the pre-granulosa enriched accessible TEs, the proximal genes were statistically enriched for sex determination related GO terms but also for terms reflecting Sertoli cell functions such as epithelium morphogenesis and angiogenesis, among other terms (**Sup. Data 5**). We also noticed that the TESCO SINE B4 observed in the pre-granulosa cells and marked with H3K27me3 is present in the H3K27ac Sertoli accessible TEs, demonstrating that TE loci are taking part in cell type-specific enhancers. Interestingly, three intergenic H3K27ac TEs were found 525 kb downstream of *Fgf9* (**Sup. Data 4**). These three TEs are contained within the 306 kb locus previously identified as an *Fgf9* enhancer-containing locus using 3D genome enhancer hub prediction tool. Deletion of the 306 kb region causes partial to complete male-to-female sex reversal (Mota-Gómez et al. 2022). Finally, the last group contains TEs showing none of the investigated histone marks (9,367 out of 31,534).

Together, these results show that TEs that gain accessibility in a time- and sex-specific manner in pre-granulosa and Sertoli cells upon sex determination are also showing promoter-like and enhancer-like properties that implies they are acting as *cis*-regulatory elements during cell differentiation. The presence of TEs within known critical sex determination enhancer loci such as TESCO, or the downstream *Fgf9* enhancer suggest the presence of other TE sequences involved in the process. Absence of any of H3K4me3 or H3K27ac on a large fraction of accessible TEs interrogates on the possibility of the presence of other histone marks, such as the recently described H4K16ac that contributes to TE transcription and *cis*-regulatory activity (Pal et al. 2023).

### Pre-granulosa and Sertoli accessible TEs are enriched in gonad transcription factor binding motifs

TE frequently contains sequences that can attract cell-specific transcription factors to promote their expression and hence their transposition. Consequently, TE transposition has dispersed transcription factor binding sites throughout mammalian genomes over evolution. Numerous studies on TEs expression in stem cells and early embryos showed that the active TE families were enriched in binding motifs for pluripotency transcription factors such as NANOG, OCT4 or SOX2. Furthermore, tissue-specific TE transcription and *cis*-regulatory element activity revealed they contain lineage-specific transcription factor binding sites (review: (Fueyo et al. 2022)). As such, we asked whether the TE loci that gain accessibility in a time- and sex-specific manner in pre-granulosa and Sertoli cells are enriched in motifs for the gonadal transcription factors, and in particular, if the motif enrichment changes depending on the chromatin landscape or the sex. We performed specific transcription factor motif enrichment analysis for known critical gonadal factors: GATA4/6 (Padua et al. 2014, Padua et al. 2015), NR5A1 (Luo et al. 1994), WT1 (Kreidberg et al. 1993), FOXL2 (Schmidt et al. 2004), RUNX1 (Nicol et al. 2019), SRY/SOX (Portnoi et al. 2018, Sinclair, et al. 1990, Wagner et al. 1994) and DMRT1 (Raymond et al. 2000). We used an approach of motif-scanning that calculates the enrichment of motif occurrence per kb compared to control sequences. We first examined if TE sequences are naturally enriched in gonadal transcription factor motifs. To this aim, we selected 3 x 10,000 random TE loci across the genome and compared their sequence composition to motifs with 3 x 10,000 random DNA regions that are not TEs (**Figure 5A**). We found that the motif recognized by NR5A1, that is involved all along gonadal differentiation in mammals was enriched in randomly selected TE loci (control TEs) from the mouse genome when compared to regions containing no repeated elements (non-TE control). For the other motifs analyzed, we did not find any significant enrichment in TE; some motifs as GATAs, WT1 and RUNX1 are even under represented in TEs.

**Figure 5.**
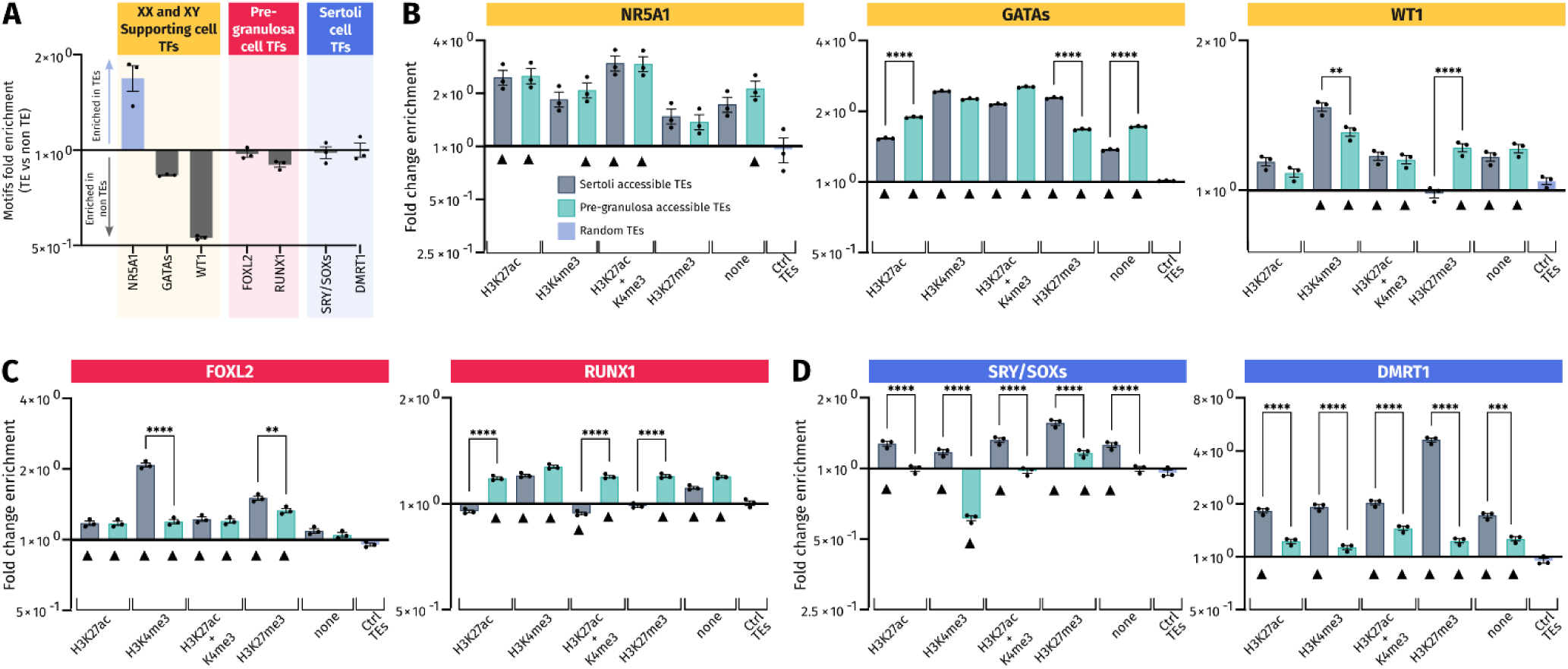
Transcription factor binding motif enrichment in pre-granulosa and Sertoli cell accessible TE loci. **A)** Ratio of the number of the respective transcription factor motifs found in 3 sets of 10,000 randomly picked TE or non-TE sequences, respectively. For each set of sequences, motifs analyzed were normalized in matches per kb and were expressed (in Log2) by the ratio of matches per kb in TE/ matches per kb in non-TE. **B-D)** Enrichment for the transcription factor motif in motifs per kb of the different analyzed sets of accessible TEs reported to the motifs per kb of three different sets of 10,000 control TEs. Black triangles mark the accessible TEs having a significant enrichment of each motif compared to control TEs (ctrl TEs) (see details of significance in **Sup. Data 6**). Motifs in **B** are those recognized by transcription factors involved in specification of XY and XX somatic progenitor cells. Motifs in **C** are those for transcription factors involved in differentiation of pre-granulosa cells, and motifs in **D**, are corresponding to Sertoli cells.

We then compared the accessible TEs harboring or not some histone marks to the control TEs randomly distributed in the genome. All the gonadal motifs examined were enriched in almost all the analyzed TE regions compared to the control TEs, regardless of the associated histone mark (**Figure 5B, C and D**). This suggests that TEs recruited during gonadal differentiation may have acquired these motifs *de novo*. It is particularly the case for the NR5A1 motif (**Figure 5B**) that is already enriched when compared to the non-TE control (**Figure 5A**).

Importantly, we also found sexual dimorphism between Sertoli and pre-granulosa in TEs bearing the same histone marks. Interestingly, motifs for male-specific transcription factors as SRY/SOXs and DMRT1 were more represented in Sertoli accessible TEs than in pre-granulosa cells independently of their associated histone marks (**Figure 5D**). Likewise, pre-granulosa specific transcriptions factors (FOXL2 and RUNX1) also displayed some sexual dimorphisms (**Figure 5C**) but to less extent than what we observed for the Sertoli specific ones. Finally, motifs for transcription factors involved in the formation of the primordial gonads, and later on the maintenance of both Sertoli and pre-granulosa cells, displayed less sexual dimorphism as GATAs, WT1, and NR5A1 which were present at the same frequency in both Sertoli and pre-granulosa cells accessible TEs (**Figure 5B**).

Our results demonstrate that accessible TEs in Sertoli and pre-granulosa cells are enriched in motifs recognized by the major transcription factors involved in gonadal differentiation. In addition, the motifs recognized by male transcription factors are globally more present in TEs opened in Sertoli cells than in pre-granulosa cells in all subsets of accessible TEs. This suggest that the accessible TEs in male and female supporting cells may have acquired *de novo* motifs recognized by transcription factors involved in the regulation of their respective transcriptional program.

## Discussion

In this work, we analyzed, for the first time, the TE ecosystem in mouse developing gonads during the time window of sex determination. We observed that around 17,000 individual TE loci are transcribed independently of gene promoters and that their expression is tightly regulated in a sex- and time-specific manner as the bipotential gonads adopt their ovarian or testicular identity. Some of the differentially expressed TE loci are found in close proximity to protein-coding genes (100 kb) and their expression correlates with their proximal genes. At the chromatin level, we showed that TE loci are acquiring cell-type specific enhancer-like and promoter-like characteristics while the gonadal somatic cells specify as pre-granulosa or Sertoli cells in XX and XY gonads, respectively. DNA motif analysis revealed that TE loci are displaying potential binding sites for the critical gonadal development and sex determining transcription factors. This study resulted in the identification of TE-derived regulatory elements that can participate and contribute to the complex gene regulatory network controlling gonadal development in mice.

Unbiased measurement of TE expression at a locus-specific level remains a challenge using shotgun sequencing technologies. Due to their repetitive nature, short reads cannot be easily attributed to a specific locus, unless the TEs have accumulated mutations along evolution. As such, despite the use of Expectation-Maximization (EM) algorithms that re-allocate multi-mapped reads, TE quantification is biased toward ancient TEs compared to recently inserted TEs displaying fewer mutations. The same challenge is even aggravated when analyzing genomic sequencing data such as ATAC-seq and ChIP-seq, as no specific tool exist to attempt to re-allocate the multi-mapped reads. Hence, the current analysis is underestimating the number of individual TE loci that are transcribed or accessible in the mouse gonads. The use of long read sequencing technologies would allow to unambiguously map individual TE loci with fragments longer than the TE themselves and would result is a more resolutive overview of the TE landscape.

Our analyses are based on RNA sequencing of whole gonads and raise the question as to whether the TEs might be expressed in a cell-type specific manner. We first intended to re-analyze single-cell transcriptomic data (Stevant, et al. 2019, Stevant et al. 2018), however such technologies revealed to not be sensitive enough to efficiently detect TE expression (10,000 total expressed TE loci in average per stage, against 75,000 in the present bulk RNA-seq, data not shown). Using publicly available bulk RNA-seq data, we could evaluate whether the expressed TE we measured in the whole gonads were originating from the somatic cells or the germline compartments. We re-analyzed the datasets obtained at E11.5 from purified somatic precursor cells, and showed that at this stage, half of the self-expressed TEs expressed in the whole gonads were commonly expressed in the purified somatic cells. Moreover, previous analysis at E13.5 (Liu, et al. 2014) suggests that some endogenous retroviruses are much more expressed in somatic than the germ cells of the fetal testes. Ultimately, high quality RNA-seq data of purified cell types composing the developing gonads of both sexes, similarly to what has been done using microarray technology (Jameson, et al. 2012), would greatly contribute to the field to study yet unexplored aspects of the gonadal transcriptome with a better sensitivity than currently existing single-cell data.

An important question is: what might be the role of these expressed TEs during gonadal development? In the early embryo, which is the most investigated model for TEs expression to date, it has been shown that activation of LINE1 expression is required for the global chromatin accessibility, independently of the coding nature of the transcript (Jachowicz, et al. 2017). While rarely present in protein-coding genes, co-opted TEs, and particularly retrotransposons, constitute a source of lncRNAs (Kapusta, et al. 2013, Kelley and Rinn 2012) known to be necessary for the correct execution of cellular processes. The endogenous MERVL, expressed in the 2-cell stage (Svoboda et al. 2004), is a marker of totipotent cells (Macfarlan et al. 2012). The TE transcript, but not the encoded protein, is required for the correct development of the pre-implantation embryo, possibly through chromatin remodeling during the totipotent-to-pluripotent transition. At the mechanistic level, it has been shown that LINE1 RNA recruits Nucleonin and TRIM28 to regulate some target genes in embryonic stem cells (Percharde, et al. 2018). During neurogenesis, TE RNAs are associated with chromatin and regulate the activity of polycomb repressor complexes PRC2 (Mangoni et al. 2023). The dynamics of TEs expression suggests that some TEs expressed uniquely at E10.5 could contribute to the multipotent state of the cells prior to sex determination, while TEs expressed during and after cell specification and in a sex-specific manner might be involved in the cell differentiation process. Targeted transcriptomic silencing or ablation of these TEs loci using CRISPR technologies would allow to functionally explore their role and understand the mechanism of action.

Chromatin architecture influences gene transcription by modulating the access of *cis*-regulatory elements to transcription factors. In this study, we identified that half of the DNA regions that gain accessibility while the gonadal progenitor cells differentiate as pre-granulosa and Sertoli cells overlap with TE loci. A large proportion of these TEs are displaying distinctive histone marks for enhancers and promoters (H3K4me3 and H3K27ac). Using a similar approach, previous studies have identified lineage specific TE-derived *cis*-regulatory elements (see review: (Fueyo, et al. 2022)). However, functional evaluation survey in mouse embryonic and trophoblast stem cells using genome editing technologies revealed that few of them are critical for gene regulatory networks (Todd et al. 2019). We predict that most of the candidate TE-derived cis-regulatory regions identified in this study are not critical players of supporting cell differentiation but rather redundant or shadow enhancers that help to sustain the gene regulatory network. However, we cannot exclude that particular TE loci might have become key players of the gonadal sex determination process. The presence of TEs within TESCO suggest their role in its enhancer activity. The TESCO sequence is highly conserved across mammals (Bagheri-Fam et al. 2010). In mice, TESCO contains four different TE sequences (one DNA TcMar-Tigger and three LINEs including a B4 and two MIR). These four TEs are conserved in rats, while only the two LINEs from the MIR (mammalian interspersed region) family are found across multiple mammalian species such as Rat, Rabbit, Human and Tree shrew (cf. https://genome.ucsc.edu). These TEs contains predicted binding motifs for SOX and GATA family of transcription factors (cf. https://genome.ucsc.edu). As such, they potentially contribute to species-specific characteristics of TESCO activity. In the same line, several studies have shown that TEs are highly heterogenous between mouse strains (Jung et al. 2023, Nellaker et al. 2012) that can modify the genes expression (Zhou et al. 2021) and chromatin dynamic (Ferraj et al. 2023). Therefore, we can speculate that strain-specific TEs could be involved in the sensitivity of C57BL/6J background to sex reversal (Eicher et al. 1996, Eicher et al. 1982, Munger et al. 2009).

The enrichment of critical gonadal transcription factor binding motifs in these accessible TE loci in the differentiating supporting cells further substantiates that TEs could have been co-opted as *cis*-regulatory elements in the context of gonadal sex determination. Enrichment of TE sequences in NR5A1 binding motifs, regardless of their chromatin landscape, shows that this motif is seemingly already present in the ancestral TE sequences, and will be even more enriched in the accessible TEs of both sexes. By contrast, specific enrichment of gonadal transcription factor motifs in the accessible TE loci in a time- or sex-specific manner could be explained by *de novo* motif acquisition and/or by co-optation of a particular TE subfamily naturally displaying these motifs (Garcia-Perez et al. 2016). To support this, our results show a clear enrichment for male-specific transcription factors (SRY/SOXs and DMRT1) for the TEs specifically accessible in Sertoli cells compared to pre-granulosa cells. A more refine study of motif enrichment in TE subfamily consensus sequences and mouse locus-specific TEs would reveal whether the pre-granulosa or Sertoli accessible TEs have evolved as gonadal-specific *cis*-regulatory elements. Again, concerning strain specificity, the motifs composition might vary depending of the mouse strain and influence subsequent recognition by DNA-binding proteins as it has been shown for CTCF in mouse CD-1 strain (Jung, et al. 2023).

In light of our finding, suggesting that TEs may participate in sex determination, it will be interesting to re-analyze human patients suffering of difference of sexual development (DSDs) with unexplained etiology (more than 50%: (Rakover et al. 2021)) to see whether reshuffle of TEs loci might be the cause for their phenotypes. Indeed, development of long-reads technologies of sequencing, and dedicated bioinformatic tools for analysis of repeated sequenced, would allow to better resolve these questions.

## Material and methods

### RNA-seq mapping and quantification

RNA-seq FastQ files from the *Trim28* knockout mouse ovaries (Rossitto, et al. 2022) (GSE166385), the *Dmrt1* ovarian reprogramming (Lindeman, et al. 2015) (GSE64960), the whole embryonic gonad splicing event (Zhao, et al. 2018) (SRP076584), and Nr5a1-GFP+ purified somatic cells at E11.5 (Miyawaki, et al. 2020) (GSE151474) studies were downloaded from Gene Expression Omnibus using the nf-core/fetchngs pipeline v1.10.0 (Ewels et al. 2020, Patel 2023a).

Gene and locus specific TE expressions were measured using SQuIRE v0.9.9.92 (Yang, et al. 2019). Briefly, FastQ files were mapped with STAR v2.5.3a on the mm10 mouse reference genome from UCSC. Gene and TE quantification were performed using the mm10 gene annotation as well as RepeatMasker TE annotation from UCSC. First, uniquely mapped reads were assigned to their corresponding TE loci. Then, the multi-mapped reads were assigned to TEs loci using an Expectation-Maximization (EM) algorithm. A score was calculated for each TE locus to account for the proportion of reads uniquely mapped and re-attributed multi-mapped reads in the total TE read count. As TEs with few uniquely aligning reads and lots of re-attributed multi-mapped reads may be prone to low confidence quantification, we considered TEs displaying a score > 95 and a minimum of 5 reads covering the TE locus. We also excluded TEs located outside the conventional chromosomes. Subsequent analysis was performed with R version 4.2.2 “Innocent and Trusting”. TE counts together with gene counts were normalized by library size using DESeq2 prior to analysis (Love et al. 2014).

### Expressed TE classification and annotation

TEs were classified as gene-dependent or self-expressed if they were found within expressed genes or in intergenic regions or non-expressed genes. To proceed, SQuIRE read counts for genes and TEs were loaded in R. TE loci and mm10 gene annotation GTF files were transformed as Genomic Ranges objects using the GenomicRanges (Lawrence et al. 2013) and the rtracklayer packages respectively (Lawrence et al. 2009). TE and gene overlaps were computed using ‘GenomicRanges::findOverlaps’ with the option ‘ignore.strand=TRUE’. For each sex and embryonic stage, we checked whether the genes overlapped by TEs are expressed with a minimum of 10 reads per gene. TEs located within an expressed gene were classified as gene-dependent as we consider their expression driven by the gene promoter, and the rest of the TEs was classified as self-expressed.

Annotation of TEs as intergenic, exonic, or intronic was performed using ‘ChIPseeker::annotatePeak’ (Wang et al. 2022) using the same mm10 GTF file as the TE quantification with SQuIRE for genome annotation version consistency.

### TE Family enrichment test

TE family enrichment test was performed using the ‘fisher.test’ R function as described in (Chang, et al. 2022). For each TE family, we built a contingency table with the number of expressed TEs belonging or not to the family and all annotated TEs belonging or not to the family and we applied the Fisher-exact test. P-values were corrected for false discovery rate using ‘p.adjust’ function with the Benjamini-Hochberg method.

### TE differential expression analysis

Differential TE expression analysis by embryonic stage and sex were performed using DESeq2 using both TE and gene quantification for a correct normalization of the data. First, a sample sheet was built with the ‘stage_sex’ sample annotation (*e.g.* ‘E10.5_XX’) for each sample. A DESeq2 object was created using ‘DESeqDataSetFromMatrix’ with ‘∼stage_sex” as design.

Sexually dimorphic TEs were obtained with the ‘DESeq’ function with default parameters (Wald test). XX and XY overexpressed genes per stage were recovered using the ‘results’ function with “stage_XX’ and ‘stage_XY’ contrast for each stage. Dynamically expressed TEs were computed sex independently. For each sex, dynamic TE were obtained using the ‘DESeq’ function with ‘test=“LRT”, reduced = ∼1’ options (Likelihood ratio test). In both differential expression analysis, we considered a TE statistically differentially expressed with an adj.pval<0.05.

Dynamically expressed TE per sex were then represented as a heatmap with complexeHeatmap (Gu et al. 2016) with rows normalized with z-scores and classified into seven clusters using ‘Ward.D2’ clustering method. The number of clusters was chosen by visual inspection of the heatmap split.

### TE-gene correlation

To inspect whether TE expression was correlating with nearby gene expression, we calculated TE-gene expression correlation per sex across the different embryonic stages. To identify the genes located near the time- and sex-differentially expressed TEs, we first checked the average distance of a TE and the nearest gene TSS. We observed that most of the TEs are located up to 100 kb from a gene TSS (**Sup. Figure 2D**). We defined a window of 100 kb upstream and downstream of the time- and sex-differentially expressed TEs and selected all the genes overlapping these windows. We then calculated the Pearson correlation between the TEs and their nearby genes expression across all embryonic stages. We considered a positive correlation with ρ>0.6 and p-value<0.05, and a negative correlation with ρ<-0.6 and p-value<0.05.

To visualize the expression correlation between TEs and nearby genes, we used the bigwig files generated with SQuIRE with the ‘draw’ script. In order to have a clean representation of the TE expression and avoid seeing other nearby TEs that were previously excluded by our quality filters described above, we used the bigwig files containing the uniquely mapped reads only and not the re-attributed reads. The genomic track visualization was built using Gviz R package (Hahne and Ivanek 2016).

### ATAC-seq mapping and analysis

FastQ files for the ATAC-seq data from E10.5 and E13.5 early gonadal progenitors and supporting cells respectively (Garcia-Moreno, Futtner, et al. 2019) were downloaded from Gene Expression Omnibus (GSE118755) using the nf-core/fetchngs pipeline v1.10.0. Data were mapped on the mm10 reference genome and analysed using the nf-core/atacseq v2.1.2 (Patel 2023b). Sequencing data quality was assessed using FastQC, sequencing adapters were removed with Cutadapt, and reads mapped using BWA. Reads were filtered with Picard to remove unmapped and duplicated reads. Peaks were called with MACS2 using the “narrow peak” parameter. For the subsequent analysis, we selected the peaks found in at least two replicates.

Differential accessibility analysis between XX and XY for each stage as well as E10.5 and E13.5 for each sex was performed using DESeq2. First, a consensus table of the read counts per accessible regions was built with featureCounts using the nf-core/atacseq pipeline. Time-specific and sexually-dimorphic accessible regions were obtained with the ‘DESeq’ function with default parameters. Sex biased accessible regions per stage and stage biased accessible regions per sex were recovered using the ‘results’ function with “stage_XX’ and ‘stage_XY’ contrast for each stage and “E10.5_sex’ and ‘E13.5_sex’ respectively. A region was defined as differentially accessible with an adj.pval<0.05 and |log2(foldChange)|>1. TE overlapping the differentially accessible regions were recovered using ‘GenomicRanges::findOverlaps’ with the option ‘ignore.strand=TRUE’.

### ChIP-seq mapping and analysis

FastQ files for the ChIP-seq data (H3K4me3, H3K27ac, H3K27me3) from E10.5 and E13.5 early gonadal progenitors and supporting cells, respectively (Garcia-Moreno, Futtner, et al. 2019) were downloaded from Gene Expression Omnibus (GSE118755, GSE130749) using the nf-core/fetchngs pipeline v1.10.0. Data were mapped on the mm10 reference genome and analysed using the nf-core/chipseq v2.0.0 (Ewels 2022). Similarly to the ATAC-seq analysis, sequencing data quality was assessed using FastQC, sequencing adapters were removed with Cutadapt, and reads mapped using BWA. Reads were filtered with Picard to remove unmapped and duplicated reads. Peaks were called with MACS2 with the respective input controls using “narrow peak” parameter for H3K4me3, and “broad peaks” for H3K27ac and H3K27me3. Because of the differences of sequencing depth, mapped reads and peak numbers between the replicates, we considered all the peaks called from any replicates for the rest of the analysis.

### TE loci epigenetic classification

We first selected the TEs that contributes to the differentiation of the supporting cells as either Sertoli or pre-granulosa cells. To proceed, for each sex separately, we selected the TE displaying an accessible chromatin region that is statistically more accessible at E13.5 than E10.5, as well as the accessible chromatin regions that are sexually dimorphic at E13.5. We extended the coordinates of the obtained TEs by 200 bp to look for ChIP-seq peaks in the direct vicinity of the TEs, at a one nucleosome resolution. We overlapped the obtained TE regions with the ChIP-seq peaks from E13.5 H3K4me3, H3K27ac and H3K27me3 for both sexes. For each TE, we binarized the presence of either H3K4me3, H3K27ac or H3K27me3, *i.e.* we marked 1 if one or more peak were found or 0 of no peak was found, and check the presence of the following combination of histone marks that are biologically relevant: only H3K4me3 (promoters), H3K4me3 and H3K27ac (promoters), only H3K27ac (active enhancer), only H3K27me3 (silencer), H3K4me3 and H3K27me3 (poised promoters). The other possible combinations were labeled as “Others”, and the loci overlapping none of these histone marks were labeled as “No peak”. The histone marks overlapping the open TEs were represented on heatmaps with EnrichedHeatmap (Gu et al. 2018) using merged replicate bigwig files produced with wiggletools and bedGraphToBigWig utilities. For visualization clarity, the combinations of histone marks with fewer than 500 open TEs were grouped in the “Other” section as the groups were not clearly visible on the final heatmap.

### GO -term enrichment analysis

Biological process GO term enrichment analysis was performed using the g:profiler website (https://biit.cs.ut.ee/gprofiler/gost) using the mouse genome as background, Benjamini-Hogberg FDR < 0.5. Because the large amount of genes present in the different analysis, the GO terms were displaying very general processes such as “Developmental processes”. For the concision of the reported results, we filtered the obtained statistically enriched GO terms to keep biologically relevant terms containing “sex”, “reproduction/reproductive”, “gonad”, but also “Wnt” in female-related gene lists, and “angiogenesis” and “epithelium” for male-related gene lists.

### TF motif enrichment analysis

Binding sites for transcription factors involved in mammalian sex determination / differentiation were counted using the Matrix scan (full options) algorithm from the RSAT suite (http://www.rsat.eu) (Santana-Garcia et al. 2022). The DNA-binding preferences for the transcription factor analyzed were modeled as matrices and analyzed with default settings and p-value set to 10^-4^. Matrices for transcription factors were obtained from JASPAR database (https://jaspar.genereg.net): SRY (MA0084.1); SOX9 (MA0077.1); SOX8 (MA0868.2); SOX10 (MA0442.2); SF1, also known as NR5A1 (MA1540.1); GATA4 (MA0482.1); GATA6 (MA1104.1); DMRT1 (MA1603.1); FOXL2 (MA1607.1); RUNX1 (MA0002.2) and WT1 (MA1627.1). GATA4 and GATA6 are redundant in gonadal differentiation (Padua, et al. 2014, Padua, et al. 2015) and the HMG domain of SOXs proteins display related function(Bergstrom et al. 2000, Polanco et al. 2010). Therefore, the pooled results of matrices scanning for GATA4 and 6 were expressed as “GATAs”, and “SRY/SOXs” for pooled scanning with SRY, SOX8/SOX9/SOX10 matrices. To normalize counts, each motif was expressed in the number of matches per kilobases of the total length of the datasets. Statistical analyses were made using GraphPad PRISM 10 with ordinary one-way Anova and Tukey’s multiple comparisons test. Detailed results of the statistical tests are provided as a supplementary table (**Sup. Data 6**).

## Supporting information

Supplementary_Material_Stevant_et_al

Sup_Data_1_Dynamic_TEs

Sup_Data_2_Sex_dimorphic_TEs

Sup_Data_3_DE_TE_gene_correlation

Sup_Data_4_TE_histone_marks_classification

Sup_Data_5_GO_terms_TE_histone_marks_classification

Sup_Data_6_TF_motifs_stats

## Author Contributions

I.S. and F.P. conceptualized the analysis. I.S. analyzed the data. I.S. and F.P. wrote the manuscript. N.G. and F.P. contributed to manuscript preparation.

## Acknowledgments

We are grateful to the genotoul bioinformatics platform Toulouse Occitanie (Bioinfo Genotoul, https://doi.org/10.15454/1.5572369328961167E12) for providing the computing resources necessary for the current study. We thank Stéphanie Le Gras from the GenomEast platform of the IGBMC in Strasbourg for her preliminary work on the TE expression analysis. We thank all the members of the Poulat’s and the Gonen’s teams for the constructive discussions and comments.

