## Supplementary_Material_Stevant_et_al for "Transposable elements acquire time- and sex-specific transcriptional and epigenetic signatures along mouse fetal gonad development"

#### Supplementary Data

**Supplementary data 1: Temporally differentially expressed self-expressed TEs.**

Results of the differential expression analysis across embryonic stages for XX and XY gonads using DESeq2. Only results with padj<0.05 are reported. The column "name" refers to the TE locus identifier which is represented as chromosome|start|end|TEsubfamily:TEfamily:TEclass|size|strand. The "baseMean" is the mean of normalized counts for all samples, "log2FoldChange" of the fold change in expression of XX vs XY normalized expression conditions transformed with log2. Positive values indicate over-expression in XX, and conversely, negative values indicate overexpression in XY. "lfcSE" columns gives the standard error of the log2FoldChange. "pvalue" is the p-values resulted from the WALD test, and "padj" is the p-value corrected for false discovery rate using Benjamini-Hochberg. The column "cluster" refers to the TE clustering presented on the heatmaps from figure 2A. XX and XY results are reported in separated tabs of the spreadsheet.

**Supplementary data 2: Sexually differentially expressed self-expressed TEs.**

Results of the differential expression analysis between XX and XY gonads for each embryonic stages using DESeq2. Only results with padj<0.05 are reported. Similarly to Sup. Data 1, the column "name" refers to the TE locus identifiers, the "baseMean" is the mean of normalized counts for all samples, the "pvalue" is the p-values resulted from the LRT test, and "padj" is the p-value corrected for false discovery rate using Benjamini-Hochberg. The columns "sex" and "stage" refer to the sex and stage in which the TE is overexpressed.

**Supplementary data 3: Sex- and time-differentially expressed TE correlation with genes in their 100 kb neighborhood.**

Results of the TE-gene correlation analysis. "TE" column refers to the TE locus identifiers, the "Gene" column are the symbol of the genes present in the 100 kb regions around the TEs. The distance of the gene TSS to the TE is referred in the "distance_to_gene" column. XX and XY Pearson correlation coefficients and their respective p-values are referred. Mathematically impossible correlation test (*i.e.* when the gene expression value is equal to 0) were annotated with a correlation coefficient of 0 and a p-value noted "NA". The "TE_coord" column gives the TE loci coordinates in a format usable in IGV genome browser for convenience.

**Supplementary data 4: Classification of the TEs gaining accessibility in a sex- or time-specific manner in pre-granulosa and Sertoli cells according to histone marks.**

The spreadsheet presents three different tabs. The first tab contains the number of accessible TEs for each different biologically relevant histone mark classification. The second and third tabs contain the classification details for the pre-granulosa and Sertoli accessible TEs respectively. The first five columns refer to the TE coordinates and their identifier, columns six to eight refers to the presence or not of the respective histone marks as binary data (*i.e.* "0" if no peak was found in the ChIP-seq data, "1" if one or more peaks were found). The "category" column refers to the classification of the TE according to the histone marks found in their sequence. "annotation" are the genomic locations of the TEs, *e.g.* exonic, intronic or intergenic. "geneid" and "distanceToTSS" refer to the nearest gene TSS and the distance from the TE loci.

**Supplementary data 5: GO-term enrichment for the proximal genes to the sex- and time-accessible TEs found in pre-granulosa and Sertoli cells according to histone marks.**

The spreadsheet presents two tabs. The first shows the results for the pre-granulosa accessible TEs, while the second the results for the Sertoli cell. GO-term enrichment results were filtered to show biologically relevant terms.

**Supplementary data 6: Details of the statistical tests used to compare the enrichments of motifs between the different sets of open TE and control TE.**

The spreadsheet presents two tabs. The first shows the summary of all statistical tests and the seconds the details of each test. $P<0.032$ (*), <0.0021 (**), <0.0002 (***), <0.0001 (****). The performed was ordinary one-way ANOVA. Multiple comparisons with no matching and pairing was performed using Tukey's test.

### Supplementary Figures

#### Sup. Figure 1

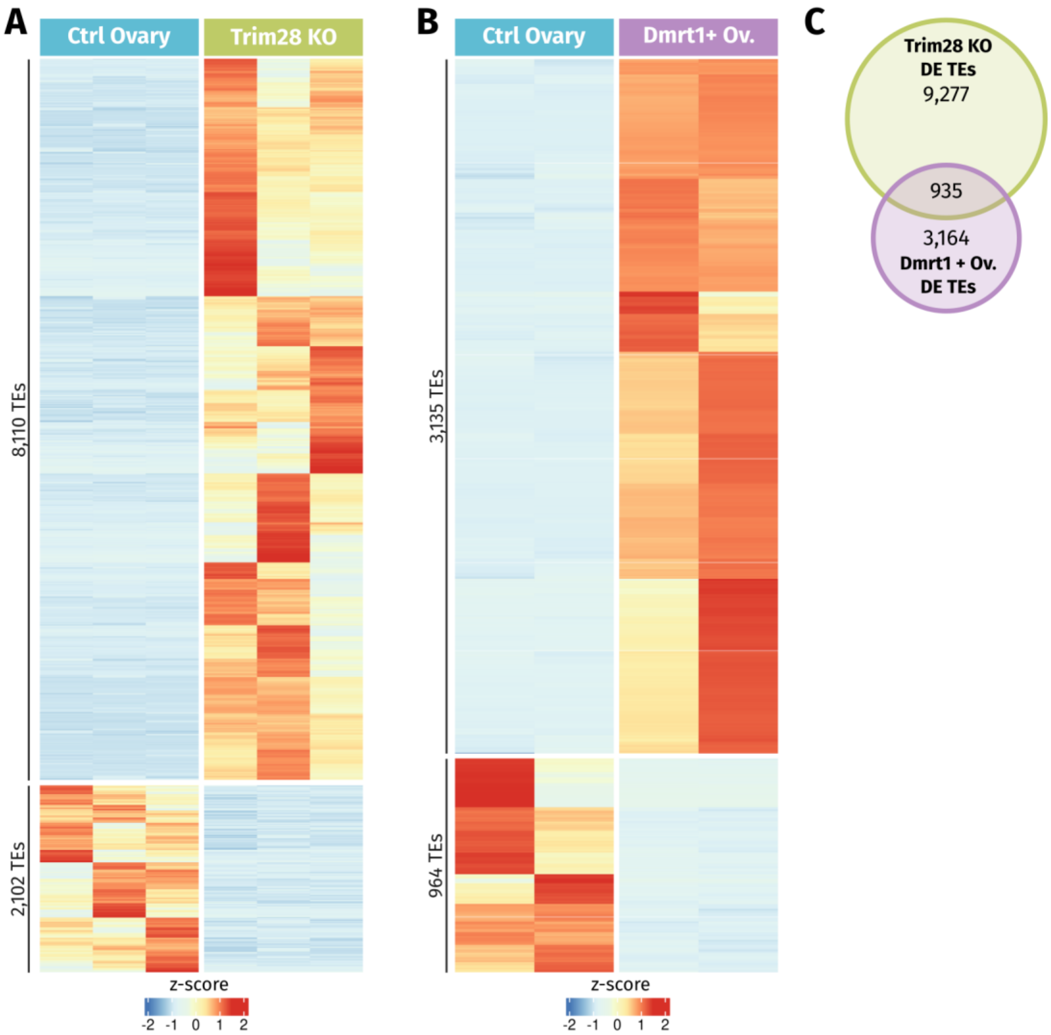


**Differential TE expression analysis of two cases of granulosa-to-Sertoli transdifferentiation in adult ovaries. A- B)** Heatmap representing the differentially expressed TEs in the *Trim28* knock-out **(A)** and the *Dmrt1* over-expression **(B)** models compared to control ovaries, respectively. Expression data were normalized using z-scores. **C)** Overlap of the differentially expressed TEs in the *Trim28* knock-out and the *Dmrt1* over-expression models. Despite the differences in mouse strain genetic background, RNA-seq library, and sequencing protocols, 935 TEs were found commonly dysregulated after granulosa-to-Sertoli transdifferentiation.

**Sup. Figure 2**

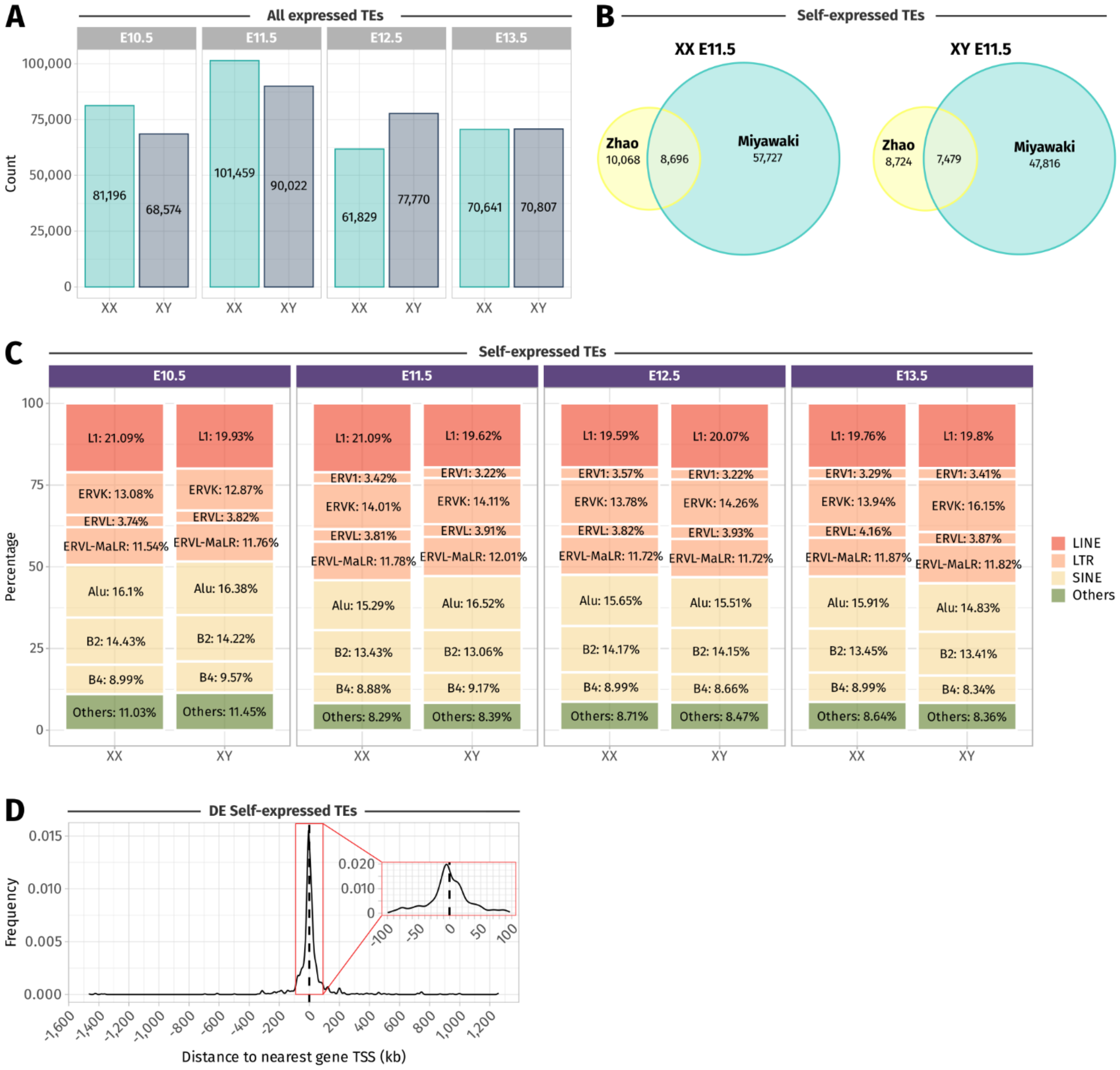


**Characteristics of the TE found expressed in mouse fetal gonads. A)** Number of TE loci detected as expressed across sexes and stages. **B)** Comparison of the self-expressed TEs detected in whole gonads (Zhao's dataset) and purified somatic cells (Miyawaki's dataset) at E11.5 in XX and XY gonads. Due to the difference in the RNA-seq library preparation protocols and sequencing sensitivity, more self-expressed are detected in the Miyawaki's dataset. However, half of the Zhao's self-expressed TEs were also found in Miyawaki's dataset. **C)** Proportion of the self-expressed TE families across sexes and stages. **D)** Distance of the sex- or time-differentially expressed self-expressed TEs relatively to the nearest gene TSS. The red square shows a magnification of the TSS region, +/- 100 kb where most of the TEs were found.

**Sup. Figure 3**

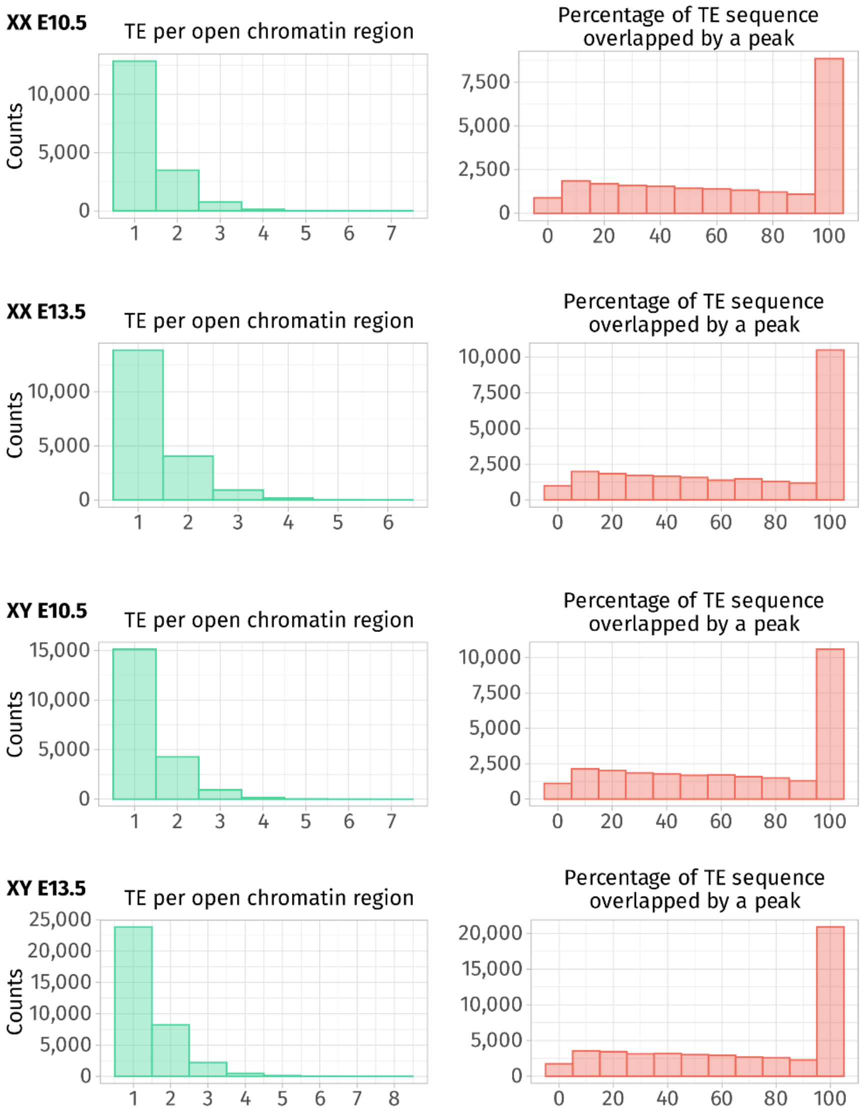


**Characteristics of the ATAC-seq peaks overlapping TEs.** Number of TEs contained within ATAC-seq peaks (green) and percentage of the TE sequences overlapping with ATAC-seq peaks (salmon) are reported for each sex and stage. The ATAC-seq peaks can contain up to eight TEs but most of them contains one or two individual TE loci. Most of the TE sequences are entirely embedded within the ATAC-seq peak (100%).
